# GBoost-CTL: A novel method in multi-tissue transcriptome-wide associations studies in cross-tissue learner incorporating GWAS information

**DOI:** 10.1101/2025.11.22.689782

**Authors:** Md Mutasim Billah, Hairong Wei, Fengzhu Sun, Kui Zhang

## Abstract

Genome-wide association studies (GWAS) have uncovered numerous genetic variants linked to complex human diseases, yet linking these variants to transcripts and tissues that drive pathology remains difficult. Multi-tissue transcriptome-wide association studies (TWAS) offer a powerful bridge, but existing analytical methods have some limitations, either by discarding important signals by separately analyzing and then aggregating results across tissues, implying imputation models in individual tissues, or fusing them with weights that ignore how much GWAS signal each tissue actually carries. Therefore, most of the existing methods do not work uniformly across different GWAS cohorts. Here, we propose GBoost-CTL - a GWAS-boosted cross-tissue learner that can overcome those aforementioned limitations. The method starts with any collection of single-tissue learners (STLs), allowing investigators to choose the most suitable imputation engine for each tissue. It then (i) allocates weights according to each STL’s out-of-sample predictive accuracy and (ii) refines those weights incorporating the GWAS-derived information, so that informative tissues are automatically up-weighted while uninformative tissues are down-weighted. This dual weighting strategy lets GBoost CTL adapt to fully shared, partially shared, or highly tissue-specific regulatory architectures while preserving nominal type I error control and delivering substantially higher power than existing linear or covariance-based methods. Through extensive simulation, we have found that this dual weighting strategy lets GBoost-CTL adapt to fully shared, partially shared, or highly tissue-specific regulatory architectures while preserving nominal type I error control and delivering substantially higher power than existing linear or covariance-based methods. When applied to real data, GBoost-CTL consistently outperformed some existing multi-tissue TWAS methods (e.g., TWAS-CTL, UTMOST and PrediXcan) by identifying a greater number of disease-associated genes with more stringent p-values. Given its modular design, computational scalability, and demonstrable gains in discovery power, we believe that GBoost-CTL offers a practical tool for the analysis of multi-tissue TWAS.

## INTRODUCTION

Genome-wide association studies (GWAS) have mapped thousands of genetic loci that affect complex human phenotypes (Tam et al., 2019; Visscher et al., 2017; Vujkovic et al., 2020), but the mechanistic path that links DNA sequence to disease remains only partially resolved. A substantial share of risk alleles act by modulating gene expression - either by altering coding sequence or through cis- and trans-acting eQTLs that tune transcriptional output (Albert & Kruglyak, 2015; Lappalainen et al., 2013; Zhang et al., 2015). Profiling gene expression at the scale needed to uncover these modest effects is often cost-prohibitive (Spencer et al., 2009), prompting the development of transcriptome-wide association studies (TWAS) (Gamazon et al., 2015; Gusev et al., 2018; Zhu et al., 2016). TWAS leverages a reference panel such as GTEx (Consortium et al., 2020): cis-SNP weights trained in the panel are used to impute expression in the GWAS cohort, and imputed expression is then tested for association with the trait. This paradigm has yielded novel biological insights across a wide array of diseases (Barbeira et al., 2018; Mancuso et al., 2017).

As multi-tissue reference resources have matured (Saha et al., 2017; Yang et al., 2017), it has become clear that modelling tissues in isolation squanders power and can even generate artefactual signals (Liu et al., 2017; Mohammadi Pejman 5 6 Park YoSon 11 Parsana Princy 12 Segrè Ayellet V. 1 Strober Benjamin J. 9 Zappala Zachary 7 8 et al., 2017; Wainberg et al., 2017). Multi-tissue approaches have enhanced the statistical power of eQTL discovery (Flutre et al., 2013; Li et al., 2018; Sul et al., 2013), indicating that a cross-tissue framework may similarly improve TWAS. UTMOST addressed part of this problem by jointly estimating weights across tissues with a sparse-group LASSO penalty (Hu et al., 2019), but it still trains each tissue independently and therefore fails to capture graded similarities among related tissues (Zhou et al., 2020). To embed homogeneity and heterogeneity directly into model building, we recently proposed TWAS-CTL, a two-stage framework that (i) trains a single-tissue learner (STL) per tissue using any predictive algorithm and (ii) fuses those predictions through an empirically derived utility weight that rewards STLs whose expression profiles generalize across tissues. Extensive simulations and an application to the GOKIND type I diabetes cohort showed that TWAS-CTL controls type I error at nominal levels and outperforms UTMOST and PrediXcan in discovery yield (Billah et al., 2025). Yet classical TWAS-CTL leaves one crucial source of information untapped. It evaluates single tissue learners (STLs) purely on cross-tissue predictive accuracy, ignoring whether the imputed expression carries GWAS signal.

Here we introduce GWAS Boosted Cross-Tissue Learner (GBoost-CTL), an augmented TWAS-CTL pipeline that incorporates both advances. For each tissue we incorporate GWAS information into the existing TWAS-CTL framework. By integrating this GWAS-level knowledge into the CTL weighting, GBoost-CTL explicitly couples imputation fidelity with downstream association potential, an idea that recent theories suggest can enhance TWAS power without inflating false positives (Bhattacharya et al., 2021; Lin et al., 2022). Through comprehensive simulations spanning multiple sample sizes, replication depths and tissue-effect scenarios we demonstrated that GBoost-CTL retained strict type I error control while delivering markedly higher power than UTMOST - particularly when heterogeneity was present in the setup. Applied to the GOKIND cohort, GBoost-CTL uncovered a richer, biologically coherent set of type-1-diabetes loci compared to TWAS-CTL, UTMOST, and PrediXcan, all at more conservative p-values than those returned by earlier methods. In short, GBoost-CTL unites cross-tissue sharing, GWAS-informed weighting and modern machine-learning prediction in a single, modular framework. The result is a next-generation multi-tissue TWAS tool that scales to the complexity of human gene regulation and, promises to sharpen our view of the molecular mechanisms that drive complex disease.

## MATERIAL AND METHODS

### Data Acquisition and Quality Control

In this project, we utilized the GTEx (Genotype-Tissue Expression) Project version 8 (v8) dataset as the reference panel for gene expression imputation. The open-access RNA-seq data includes expression profiles from 49 human tissues across 838 donors, the majority of whom were European American (85.3%), followed by African American (12.3%), Asian American (1.4%), and Hispanic or Latino (1.9%) individuals. The cohort comprised 557 males (66.4%) and 281 females (33.5%) (Consortium et al., 2020). While previous cross-tissue TWAS implementations such as UTMOST employed RPKM (v6) or TPM (v7) normalized expression data, these methods are primarily suitable for within-sample normalization and may bias downstream analyses in cross-tissue models. Between-sample normalization methods, particularly count adjustment using upper quartile factors (CUF) and trimmed mean of M values (CTF), offer greater consistency across tissues and samples (Johnson & Krishnan, 2022). Accordingly, we downloaded raw gene read counts for each tissue from GTEx v8 and applied CUF-based normalization to enable robust cross-tissue modeling. To correct for potential confounding effects, we again refined the normalized expression data for sex, sequencing platform, the top three genotype principal components (PCs), and the top PEER (Probabilistic Estimation of Expression Residuals) factors. Access to the protected GTEx genotype dataset was obtained via dbGaP under study accession phs000424.v8.p2 (https://www.ncbi.nlm.nih.gov/projects/gap/cgi-bin/study.cgi?study_id=phs000424.v8.p2), following submission by Dr. Kui Zhang. For genotype quality control (QC), we employed PLINK v1.9 ((Purcell et al., 2007); https://www.cog-genomics.org/plink/). SNPs with a missing call rate >10% were excluded (Marees et al., 2018). Additionally, variants failing the Hardy-Weinberg equilibrium (Kalogeropoulou et al.) test (p-value < 0.05) were removed to mitigate technical and population stratification artifacts (Hosking et al., 2004). Samples with >10% missing genotype data were also discarded (Zhou et al., 2020). Individual heterozygosity rates were used to detect potential contamination or inbreeding. We excluded five samples with the highest and five with the lowest heterozygosity, following the recommendation to remove outliers exceeding ±3 standard deviations from the mean (Marees et al., 2018). This step was executed using PLINK and Git Bash (https://git-scm.com/downloads). To reduce redundancy due to linkage disequilibrium (LD), we performed LD pruning using an r^2^ threshold of 0.9, retaining relatively independent variants (Zhou et al., 2020). Moreover, SNPs with minor allele frequency (MAF) below 0.05 were filtered out to prevent spurious associations, as low-frequency variants are more susceptible to sampling noise and less likely to exhibit reliable trait associations in modest-sized cohorts (Zhou et al., 2020). After applying all QC filters - missingness, HWE, LD pruning, heterozygosity outlier removal, and MAF thresholding - the final genotype dataset included 2,645,120 SNPs across 828 individuals and was used for downstream multi-tissue TWAS analyses.

## TWAS

**Figure 1** illustrates the contemporary TWAS pipeline into a two-tier framework that traces a path from local genetic variation to disease risk. The process begins with a reference panel - the GTEx resource is a typical example - where matched genotype data (*M* SNPs in *n* donors) and RNA-seq measurements are available for dozens of tissues. Within each tissue, we train a predictive model that uses the surrounding cis-SNPs to predict gene expression levels; the resulting weight vectors capture that tissue’s regulatory architecture. Those weights are then transferred to a generally far larger GWAS cohort (*M* SNPs, *n*_1_ participants). Applying them to every individual generates tissue-specific vectors of imputed expression even though no transcriptomic data were collected. Because the same SNP predictors are reused across organs, the regulatory patterns learned in GTEx are faithfully projected onto the disease sample. In the final step, the GWAS genotypes and their corresponding predicted transcriptomes are analyzed jointly: the phenotype is regressed on each gene’s imputed expression (in one or many tissues), yielding a set of gene-trait association statistics. Taken together, the three stages-learning cis effects in a reference panel, broadcasting those effects into a GWAS panel, and testing the resulting expression–trait links - constitute the core of multi-tissue TWAS, offering a principled route to identify both the genes and the tissues most likely to mediate complex-disease risk.

**Figure 1.**
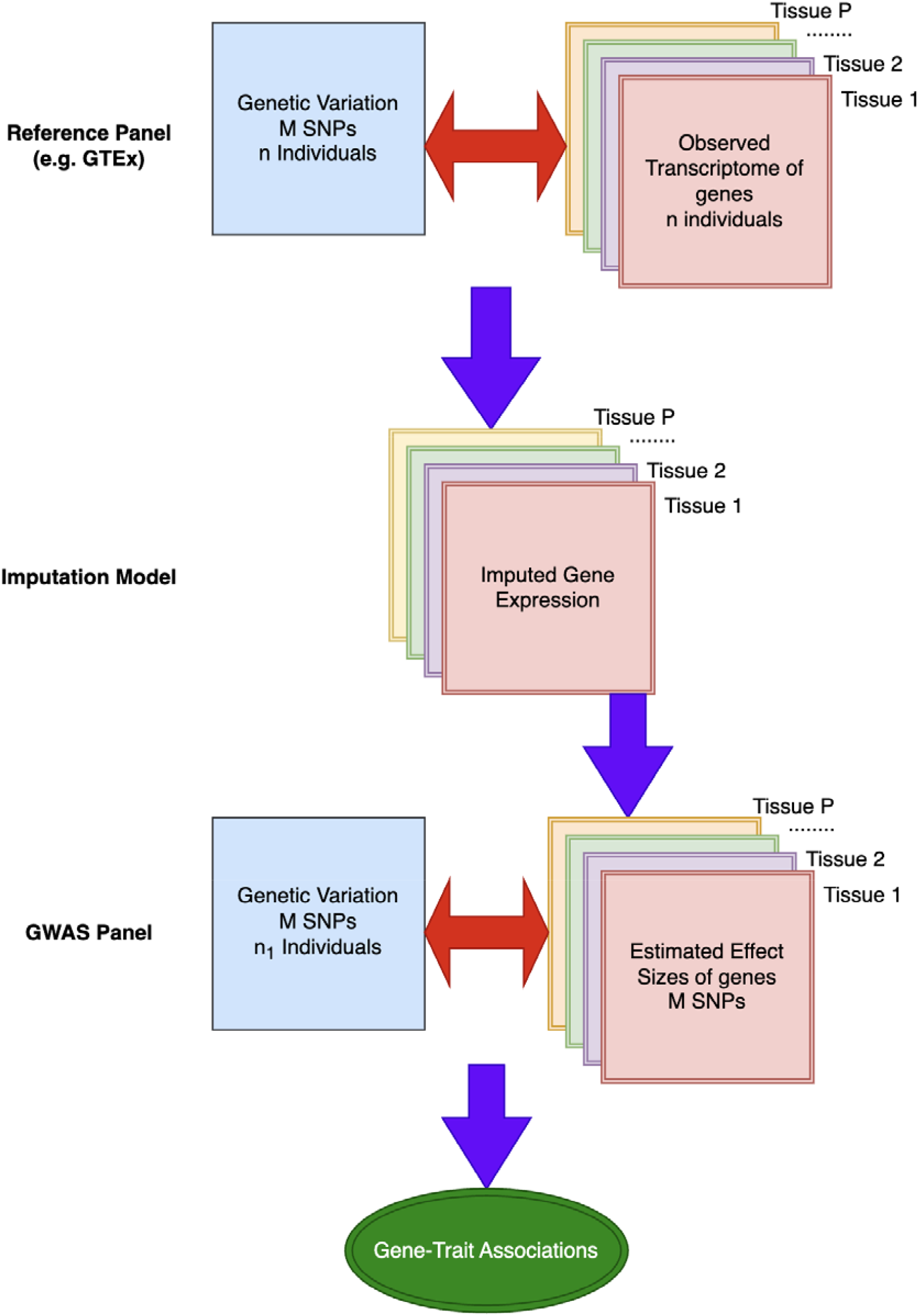
An overview of Transcriptome-wide Association Studies (TWAS).

### G-Boost-CTL

**Figure 2** (created with https://app.diagrams.net/) outlines the workflow of our proposed G-Boosted Cross-Tissue Learner (G-Boost-CTL) strategy. Starting with any target gene, we collect genotype data and gene expression measurements of different tissues from the reference panel (e.g. GTEx). In phase I, we fit a separate Single-Tissue Learner (STL) for each tissue - these learners can be penalized regression methods, tree-based regressors, deep networks, or any other methods capable of mapping cis-SNPs to expression (Billah et al., 2025). Each STL yields a tissue-specific vector of predicted expression values. Phase II, we assign reliability scores to every tissue–model pair. A bespoke weighting function up-weights predictions that generalize well and down-weights noisy or weakly predictive signals (Billah et al., 2025; Patil & Parmigiani, 2018). To harmonize the model with the downstream GWAS, in phase III, we refine the raw STL weights by the normalized variance of the predicted expression in the GWAS genotypes, so that tissues explaining more cross-individual variation receive the reasonable contribution in the adjusted weighing scheme. In the final phase (phase IV), G-Boost-CTL forms a composite, cross-tissue expression estimate by summing the STL outputs according to their adjusted weights, thereby capturing both shared and tissue-unique regulatory signals, yielding more accurate, flexible, and robust expression imputation than relying on a single tissue model or cross-tissue learner approach as mentioned in TWAS-CTL (Billah et al., 2025).

**Figure 2.**
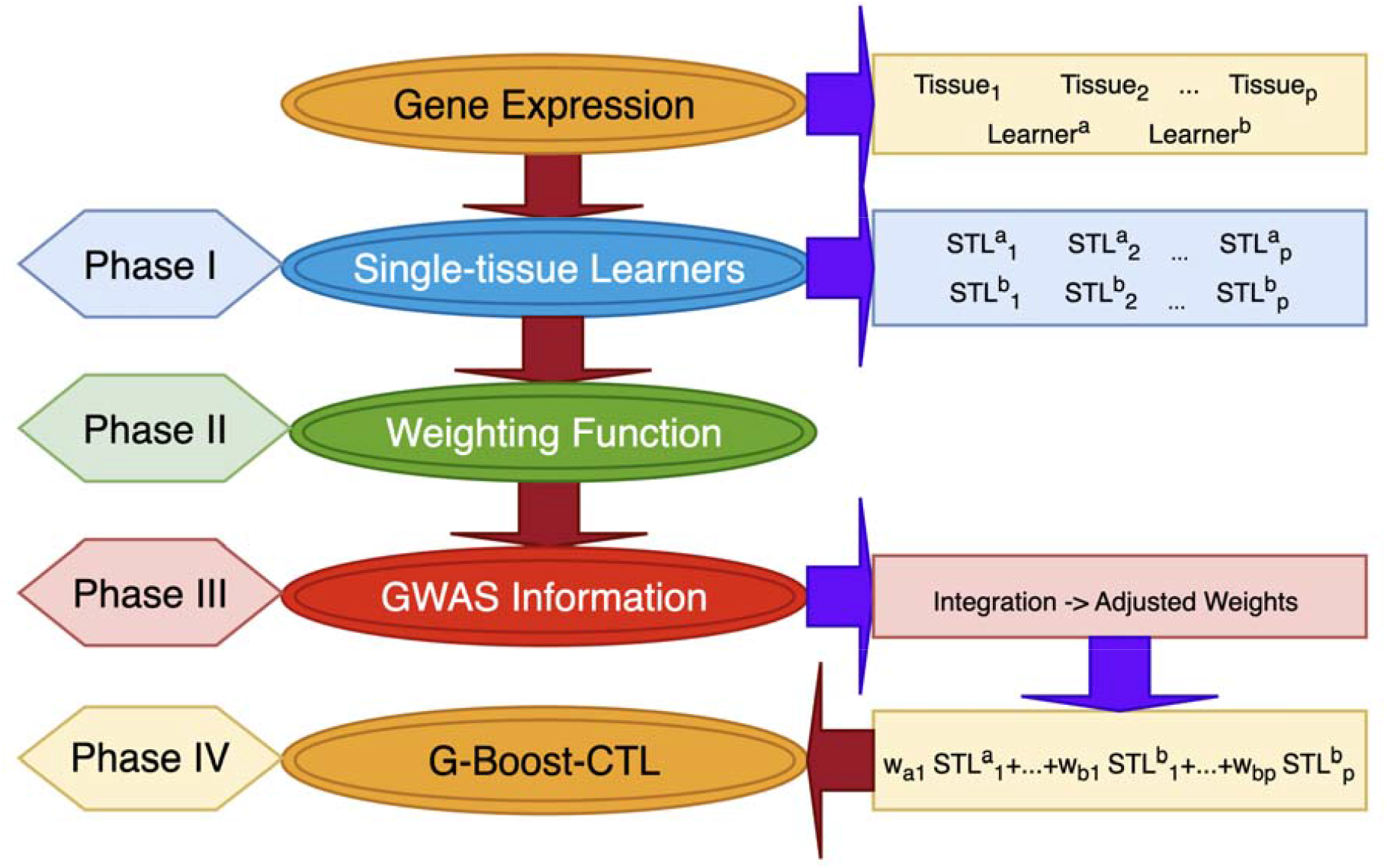
Architecture of GBoost-CTL (GWAS boosted cross-tissue learner).

For each gene, we start with genotype - expression pairs from a reference panel such as GTEx and fit an independent single-tissue learner (STL) in every tissue *j*(*j* = 1,2,…,.*P*) with learner type *l*(*l* = 1,2,…,*L*):

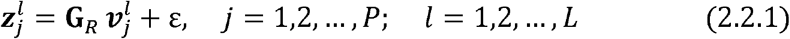

where G_***R***_ is the n × M genotype matrix in the reference panel and 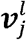 is the vector of SNP weights estimated by learner *l* for tissue *j*.

Applying those weights to the GWAS genotypes G_*w*_ (*n*_1_ ×*M*) yields the tissue-specific, imputed expression:

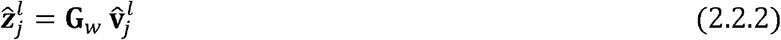

To construct a cross-tissue prediction, we aggregate the STL outputs with data-driven weights:

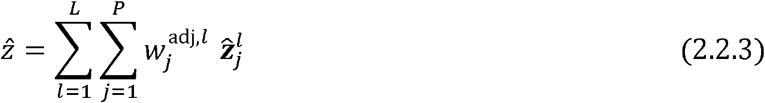

Here 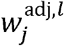 is a GWAS boosted CTL weight derived from equation (2.2.9) and the corresponding tissue-specific prediction 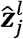 is obtained from equation (2.2.2).

To obtain 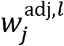, first, cross-tissue predictive performance is quantified by a utility function: 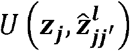, where 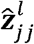 includes the predictions obtained using model 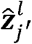, trained in tissue *j′*, on the prediction profiles in tissue *j*, using learner *l*.

For each learner, we form a *P*×*P*matrix:

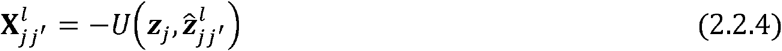

so that lower values correspond to better cross-tissue performance; each row of **X**^*l*^ therefore summarizes how well the predictor trained in tissue transports to every other tissue. For learner *l* we reduce row *j* of **X**^*l*^ to a single score by averaging the off-diagonal entries (the within-tissue resubstituting error 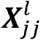 is excluded).

For each learner *l*, we derive its weights by first summarizing the row *j* of *X* by the mean of off-diagonal entries (omitting the resubstituting error 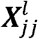):

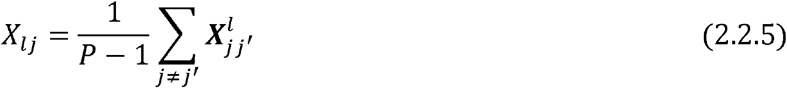

Other summaries - e.g., the root-mean of off-diagonals - can be used in place of the simple mean. The CTL weights then can be found as,

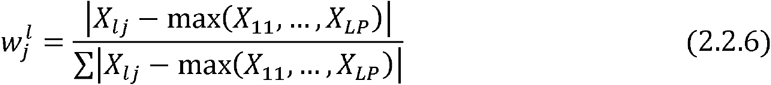

In this approach, the worst-performing STL receives a weight of zero, the losses of all other STLs are rescaled with respect to that baseline, and the adjusted values are subsequently normalized to obtain the final weights.

GBoost-CTL has the flexibility to incorporate any GWAS information as a tissue-specific regulatory weight into its framework. In this work we employ the normalized variance of the imputed expression across the GWAS cohort to quantify how much genetically driven signal each tissue contributes.

Let,

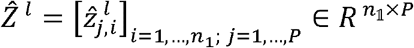

denote the matrix of predicted expression values obtained from learner *l*, where *n*_1_ is the number of GWAS individuals and the *j*-th column corresponds to tissue *j*. For tissue *j* we compute the GWAS variance:

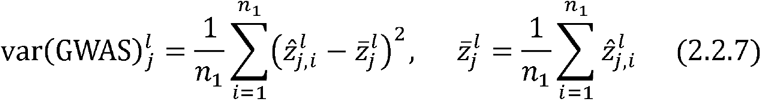

Collecting these variances into the vector 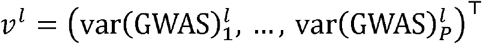, we normalize them to obtain a GWAS regulatory-weight vector:

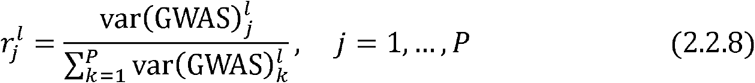

The elements of 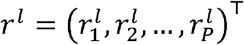 form a probability simplex 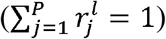 and reflect each tissue’s relative contribution to genetically regulated expression variability in the GWAS cohort.

Finally, we adjust the CTL weights obtained in Equation (2.2.6) by these regulatory factors:

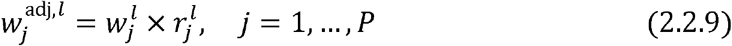

The vector 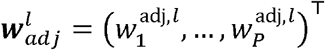 simultaneously captures (i) predictive fidelity of each STL and the degree to which the corresponding tissue’s expression varies in the GWAS sample. Empirically, this dual weighting yields a more informative cross-tissue ensemble, particularly when regulatory influence varies markedly across tissues.

As the utility function mentioned in Equation (2.2.4), For every learner and tissue pair (*j*, j ′) with *j* ≠ *j ′*, we quantify out-of-sample performance by the coefficient of determination:

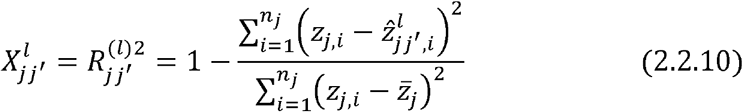

where *z*_*j,i*_ is the observed expression of individual *i* in tissue *j*,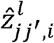 is the value predicted by the model trained in tissue *j′*, and 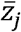 is the sample mean of tissue *j*. This choice rewards models that explain a larger fraction of the target-tissue variance. For the row-wise summary for weighting, for each learner *l* we condense the off-diagonal entries of the *j*-th row of *X*^*l*^ by an arithmetic mean, as mentioned in equation (2.2.5), as because higher *R*^2^ values denote better performance, no square-root or sign adjustment is necessary; the subsequent normalization steps (Equations 2.2.6 - 2.2.9) converts these scores into the final G-Boost-CTL weights.

### STLs: Modeling Cross-Tissue Expression Imputation

To estimate the effect size 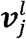 in Equation (2.2.1), any predictive modeling algorithm can serve as a candidate learner. In this study, we employ a diverse ensemble of four modeling strategies to construct our GBoost-CTL framework. Specifically, we incorporate penalized linear models - Ridge Regression (Hoerl & Kennard, 1970), LASSO (Least Absolute Shrinkage and Selection Operator; (Tibshirani, 1996)), and Elastic Net (Zou & Hastie, 2005) - which are widely used in high-dimensional settings for their regularization properties. Additionally, consistent with prior work in the field (Gamazon et al., 2015), we integrate Random Forest (Breiman, 2001), a nonparametric ensemble learning technique known for capturing non-linear interactions. Model training and hyperparameter tuning for each learner were conducted using five-fold cross-validation to ensure robust estimation and avoid overfitting.

### Gene-level Association Testing

For each gene g, we assess its relationship with the phenotype by testing the null hypothesis *H*_0_:β_*g*_ =0 against the alternative *H*_1_: β_*g*_ ≠ 0. The phenotype for every GWAS individual is described by the linear model

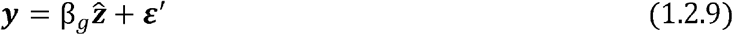

where, *y* is the trait vector, β_*g*_ is the total effect, 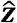 is the cross-tissue combined imputed gene expression, and 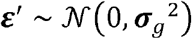.

### Simulation Architecture

To benchmark the proposed GBoost-CTL framework we drew on the Genetics of Kidneys in Diabetes (GOKIND) study (dbGaP accession phs000018.v2.p1), using both its phenotype data for real-world association tests and its genotype panel as the basis for our simulations. Access to the protected dataset was obtained via dbGaP (https://www.ncbi.nlm.nih.gov/projects/gap/cgi-bin/study.cgi?study_id=phs000018.v2.p1), and all marker coordinates were updated from their original GRCh36 positions to the current GRCh38 assembly with the UCSC LiftOver utility (https://genome.ucsc.edu/cgi_bin/hgLiftOver), ensuring compatibility with modern annotation resources. Genotypes then underwent the same quality-control pipeline applied to our GTEx reference panel - filtering on call-rate, Hardy–Weinberg equilibrium, heterozygosity outliers and minor-allele frequency. SNPs were mapped to genes through the Bioconductor biomaRt interface (Durinck et al., 2009) using GRCh37 identifiers, adopting a ±100 kb cis window motivated by GTEx v8, where more than 70 % of significant cis-eQTLs fall within this range (Consortium et al., 2020).

Our simulation strategy largely mirrors the design of UTMOST (Hu et al., 2019) but advances its ad-hoc tissue choices with a biologically motivated setup trio chosen to span both within-organ similarity and cross-organ diversity as mentioned in (Billah et al., 2025) (Nucleus Accumbens (n = 198), Brain Cortex (n = 204), and Esophagus Mucosa (n = 492)). For any specific gene, the predicted expression levels in each tissue 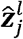 is generated by one of several single-tissue learners - Lasso, Ridge, Random Forest, Elastic-Net, and UTMOST - thereby allowing the downstream cross-tissue learners to be assessed under a variety of expression-prediction architectures. Phenotypes were then simulated as:

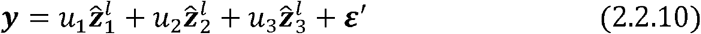

with 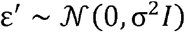 Setting all 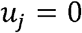 yields null data for assessing type-I error; power was examined under three regulatory regimes: (i) a homogeneous scenario (*u*_1_ = 1, *u*_2_ = *u*_3_ = 0); moderate heterogeneity (*u*_1_ = *u*_2_ = 1, *u*_3_ = 0)and (iii) full heterogeneity 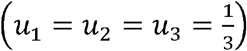. Genotype data from the GOKIND cohort were served as G_*w*_ (equation 2.2.2) when constructing imputed expression, ensuring that linkage structure and allele frequencies match those encountered in the association analysis. Each configuration was evaluated at sample sizes *n* = 500 and *n* = 750, with 1000, 5000, and 10000 Monte-Carlo replications per gene. Cross-tissue association statistics were computed for both G-Boost-CTL and the UTMOST pipeline, and power was defined as the proportion of replications achieving nominal significance at α = 0.05. This large-scale design allows a comprehensive comparison of error control and sensitivity across a spectrum of tissue-effect patterns and predictive-model pairings.

## RESULTS

### Simulation Studies: Empirical Type I Error Rate

We compared the simulation outcomes of our proposed method against UTMOST, following its implementation as detailed by (Hu et al., 2019; Zhou et al., 2020). Additionally, we incorporated PrediXcan (Gamazon et al., 2015) to impute 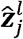 enabling a direct comparison with UTMOST. For this, we applied the Generalized Berk-Jones (GBJ) test (Sun & Lin, 2017) to integrate the imputed gene expressions across multiple tissues, consistent with the UTMOST framework. **Table 1** summarizes the empirical type I error rates obtained from G-Boost-CTL ensemble (averaged over twelve component STLs) and from UTMOST across three cohort sizes (N = 500, 750, 1000) and three Monte-Carlo replication depths (M = 1000, 5000, 10000). In every scenario both procedures remained close to the nominal 5 % level, confirming that neither approach is prone to systematic inflation. At the smallest sample size (N = 500) UTMOST displayed a noticeable upward bias when only 1,000 replicates were generated (6.2 %), whereas G-Boost-CTL tracked the target rate more closely (5.3 %). Increasing the number of replicates immediately narrowed this gap; by M = 5,000 the ensemble became slightly conservative (4.97 %), and by M = 10,000 the two methods were essentially indistinguishable. For the intermediate cohort (N = 750) the pattern reversed at low replication: G-Boost-CTL edged above the benchmark (5.35 % versus 4.80 %), yet the difference vanished once M reached 5,000, where both procedures settled just under 5 %. At the largest sample (N = 1,000) UTMOST achieved the minimum error with 1000 replicates, but the ensemble converged rapidly; by M = 10,000 G-Boost-CTL produced the lowest empirical rate (4.78 %), while UTMOST remained marginally higher (4.85 %). Taken together, these results indicate that the GWAS-boosted ensemble offers dependable control of false positives across a wide range of study sizes. Any mild inflation observed at the smallest replication depth disappears as soon as the Monte-Carlo sample is enlarged, and in the high-replication setting the ensemble is either comparable to or more conservative than the UTMOST benchmark. The stability of the error rate across ten CTLs suggests that the weighting scheme effectively shields the global test from individual learner misspecification, reinforcing the robustness of G-Boost-CTL for large-scale multi-tissue TWAS inference.

**Table 1.** Type I Error Comparison: Average type I error rate of GBoost-CTL across 12 CTLs with UTMOST.

| $N$ | $M$ | GBoost-CTL | UTMOST |
| --- | --- | --- | --- |
| **500** | 1000 | 0.0527 | 0.0620 |
|  | 5000 | 0.0497 | 0.0518 |
|  | 10000 | 0.0508 | 0.0519 |
| **750** | 1000 | 0.0535 | 0.0480 |
|  | 5000 | 0.0480 | 0.0482 |
|  | 10000 | 0.0490 | 0.0491 |
| **1000** | 1000 | 0.0507 | 0.0460 |
|  | 5000 | 0.0508 | 0.0522 |
|  | 10000 | 0.0478 | 0.0485 |

### Simulation Studies: Statistical Power

To evaluate the statistical power of our proposed G-Boost-CTL against the widely used UTMOST method, we considered three distinct cross-tissue regulatory scenarios: homogeneity, moderate heterogeneity, and full heterogeneity. For each scenario (denoted as U=1, 2, 3 respectively), we tested twelve GBoost-CTL combinations, each incorporating GWAS regulatory weights derived from a designated learner (Lasso, Ridge, Elastic Net, or Random Forest), paired with a secondary learner for association analysis. Power was estimated across 10,000 replicates using a sample size of 750.

Table 2 and **Figure 3** depicts the power comparison of different GBoost-CTL with UTMOST when the predicted gene expression was imputed using PrediXcan. When the trait was predominantly driven by a single tissue (homogeneity), methods that incorporated Random Forest (RF) for either GWAS weighting or downstream association consistently outperformed all others. Notably, the Lasso+RF (0.923), Elastic Net+Ridge (0.837), and Elastic Net+RF (0.719) CTLs achieved the highest power. These combinations exhibited an order of magnitude improvement over UTMOST (0.148), reaffirming the value of nonlinear modeling in settings with strong single-tissue influence. Even linear models like Lasso+Ridge (Ri) (0.648) and Elastic Net+Ridge (0.837) demonstrated marked improvement when paired with RF weighting. In contrast, the baseline UTMOST struggled to detect associations, indicating its limitations under pure homogeneity. In the intermediate regime of moderate homogeneity, where two of the three tissues contributed to the phenotype, nearly all GBoost-CTL combinations reached near-perfect sensitivity. Six methods achieved power ≥ 0.99, including Lasso+RF (RF), Elastic Net+Ridge, and Elastic Net+RF. Interestingly, both Ridge+RF variants (0.821, 0.927) performed exceptionally well, emphasizing the synergistic effect of combining nonlinear GWAS-informed weights with stable linear associations. UTMOST, by comparison, delivered poor performance (0.086), suggesting an inability to fully integrate partially shared signals. These results aligned with the observations from Project 1, where CTLs that could flexibly modulate tissue contribution had greater success across moderate heterogeneity. When expression-trait influence was equally distributed across all tissues, performance declines for all methods, as expected. However, GBoost-CTL models maintained a consistent lead over UTMOST. The Elastic Net+Ridge (0.370), Elastic Net+RF (0.357), and Lasso+RF (0.451) combinations achieved the highest power, nearly tripling that of UTMOST (0.102). While no method reached high absolute sensitivity in this setting, G-Boost-CTLs still captured the distributed signal more effectively. Linear–linear combinations (e.g., Lasso+Ridge (L), 0.112) underperformed, underscoring the need for flexible learners in highly heterogeneous contexts.

**Table 2.** Power Comparison of different GBoost-CTLs with UTMOST in different U settings (sample size 750, replication 10000)–when PrediXcan was used to impute 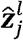. The highest power across different heterogeneity levels is bold.

| U | L+Ri (L) | L+Ri (Ri) | L+RF (L) | L+RF (RF) | R+RF (Ri) | R+RF (RF) | E+RF (E) | E+RF (RF) | L+E (L) | L+E (E) | E+Ri (E) | E+Ri (Ri) | UT |
| --- | --- | --- | --- | --- | --- | --- | --- | --- | --- | --- | --- | --- | --- |
| 1 | 0.112 | 0.648 | **0.923** | 0.098 | 0.301 | 0.221 | 0.719 | 0.240 | 0.302 | 0.684 | 0.837 | 0.077 | 0.148 |
| 2 | 0.522 | 0.999 | **1.000** | 0.044 | 0.821 | 0.927 | 0.996 | 0.913 | 0.557 | 0.983 | 0.999 | 0.038 | 0.086 |
| 3 | 0.112 | 0.438 | **0.451** | 0.050 | 0.165 | 0.212 | 0.357 | 0.197 | 0.119 | 0.287 | 0.370 | 0.049 | 0.102 |

**Figure 3.**
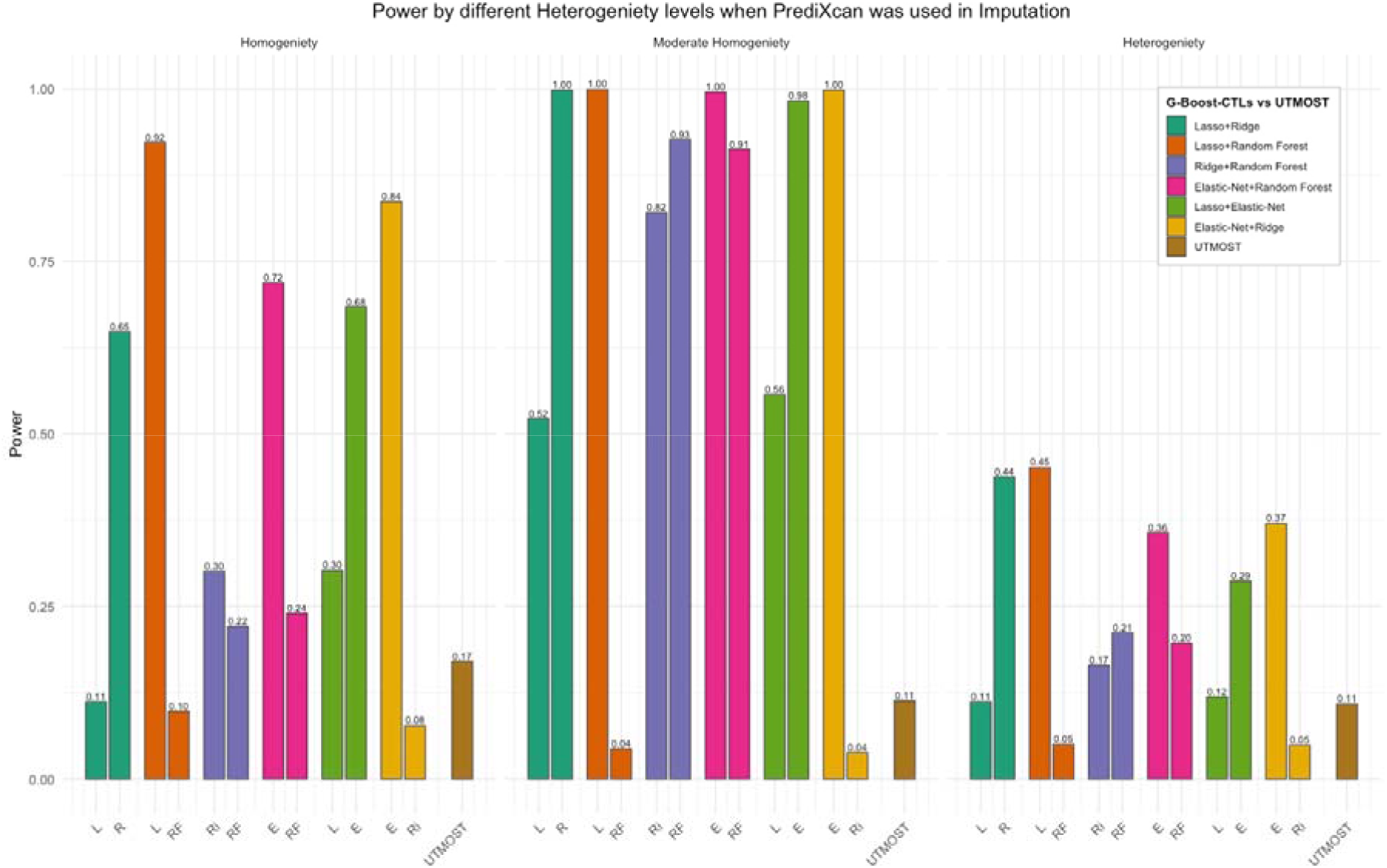
Statistical power comparison across varying levels of heterogeneity when Elastic Net is used for imputation in GBoost-CTL and UTMOST frameworks. Each panel corresponds to a different heterogeneity level (Homogeneity, Moderate Homogeneity, and Heterogeneity). Within each panel, the x-axis indicates which single-tissue learner (STL) was used to integrate GWAS information into the G-Boost-CTLs.

**Table 3 and Figure 4** illustrate the power comparison between various GBoost-CTL methods and UTMOST, based on gene expression imputed using Ridge Regression. In settings where a single tissue dominated the genetic effect, GBoost-CTL methods - especially those leveraging Ridge as the GWAS integrator - exhibited exceptional power. The highest power was achieved by combinations such as L + RF (L) and L + RF (R), both yielding 0.995, followed closely by E + Ri (E) at 0.974. These CTLs consistently outperformed UTMOST, which recorded a power of only 0.149. Interestingly, the Lasso+Ridge and ElasticNet+Ridge configurations performed robustly regardless of which learner was used to encode the GWAS feature, underscoring their strength in homogeneous settings. Under moderate heterogeneity (U=2), all G-Boost-CTLs maintained strong or near-perfect power, especially combinations with Ridge or Elastic Net in either position. For instance, L + Ri (R), L + RF (RF), and both E + RF variants all achieved power between 0.994 and 1.000. UTMOST also performed well here (0.398) but still fell short of the best G-Boost-CTLs. The E + Ri (Ri) configuration had slightly lower power (0.030), but this was a rare exception. When the genetic effect was uniformly spread across all three tissues, power declined overall, but GBoost-CTLs still held notable advantage. L + Ri (R) and L + RF (L) reached 0.770 and 0.693, respectively, outperforming UTMOST (0.178) and other less effective CTLs such as E + Ri (Ri) (0.048). Notably, Ridge-based learners integrated with Random Forest or Elastic Net consistently provided higher power under this complex, heterogeneous setting. The figure and table clearly indicate that the learner chosen to encode GWAS information (X) played a crucial role in the model’s effectiveness. For example: L + RF (L) outperformed L + RF (RF) in most scenarios, suggesting that Lasso captures GWAS-informed variability better in this pairing. Similarly, E + RF (E) consistently achieved higher power than E + RF (RF). This pattern reflected a broader trend where penalized regression learners (Lasso, ElasticNet) tended to provide more stable integration of GWAS information than tree-based learners when placed in the (X) position.

**Table 3.** Power Comparison of different GBoost-CTLs with UTMOST in different U settings (sample size 750, replication 10000)–when Ridge Regression was used to impute 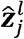. The highest power across different heterogeneity levels is bold.

| $U$ | L+Ri (L) | L+Ri (Ri) | L+RF (L) | L+RF (RF) | R+RF (Ri) | R+RF (RF) | E+RF (E) | E+RF (RF) | L+E (L) | L+E (E) | E+Ri (E) | E+Ri (Ri) | UT |
| --- | --- | --- | --- | --- | --- | --- | --- | --- | --- | --- | --- | --- | --- |
| 1 | 0.542 | **0.995** | **0.995** | 0.129 | 0.477 | 0.345 | 0.919 | 0.373 | 0.479 | 0.896 | 0.974 | 0.093 | 0.149 |
| 2 | 0.920 | **1.000** | **1.000** | 0.040 | 0.969 | 0.994 | 1.000 | 0.991 | 0.787 | 1.000 | 1.000 | 0.030 | 0.398 |
| 3 | 0.225 | **0.770** | 0.693 | 0.051 | 0.268 | 0.331 | 0.562 | 0.315 | 0.170 | 0.457 | 0.587 | 0.048 | 0.178 |

**Figure 4.**
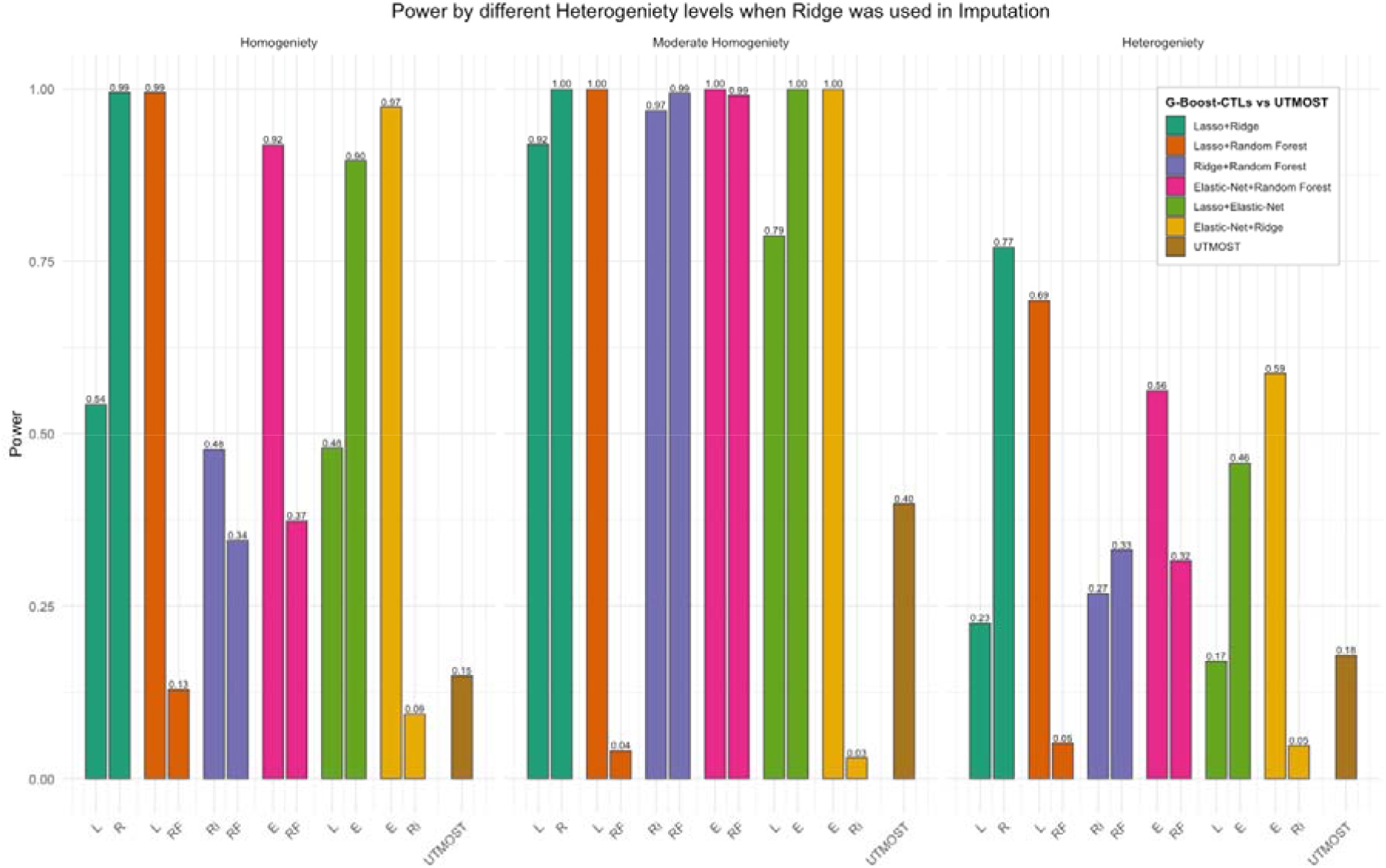
Statistical power comparison across varying levels of heterogeneity when Ridge regression is used for imputation in GBoost-CTL and UTMOST frameworks. Each panel corresponds to a different heterogeneity level (Homogeneity, Moderate Homogeneity, and Heterogeneity). Within each panel, the x-axis indicates which single-tissue learner (STL) was used to integrate GWAS information into the G-Boost-CTLs.

Table 4 and **Figure 5** depicts the power comparison between various G-Boost-CTL methods and UTMOST, based on gene expression imputation using Lasso. Under homogeneity, the best-performing GBoost CTL was L + Ri (R) with a power of 0.461, followed by L + RF (L) at 0.318 and E + Ri (E) at 0.377. UTMOST yielded a comparable power of 0.293, outperforming several CTL variants. However, most GBoost CTLs, particularly those integrating Lasso as GWAS feature (X), underperformed under homogeneity, likely due to Lasso’s sparsity-induced instability in low-variability environments. GBoost CTLs clearly outshined UTMOST in moderate homogeneity. L + Ri (R) attained near-perfect power (0.998), and several other configurations - like E+RF (E) (0.958), E + RF (RF) (0.924), and L + RF (RF) (0.672) - also demonstrated strong performance. In contrast, UTMOST recorded only 0.022, revealing a sharp drop in power under this heterogeneity level. These findings show that GBoost CTLs effectively capture partially shared tissue signals, especially when Ridge or Elastic Net is used as the boosting learner. Power dropped across the board under the configuration of heterogeneity, as expected due to the uniform spread of genetic signal across all tissues. However, GBoost CTLs like E + RF (E) (0.249) and E + RF (RF) (0.210) still outperformed UTMOST (0.107) and other combinations. This suggests that elastic net-boosted CTLs may have better resilience in complex genetic architectures compared to traditional approaches. The **Table 4** and **Figure 5** also underscore the importance of choosing the right learner for incorporating GWAS variability. As for example, L+RF (L) consistently outperformed L+RF (RF) across all U settings, particularly under moderate heterogeneity (0.540 vs. 0.639), indicating that Lasso better captured regulatory influence in this pairing. E+RF (E) outperformed its RF-integrated counterpart across all U settings, especially under U=2 and U=3, reaffirming the advantage of using regularized learners like Elastic Net for GWAS weight encoding.

**Figure 5.**
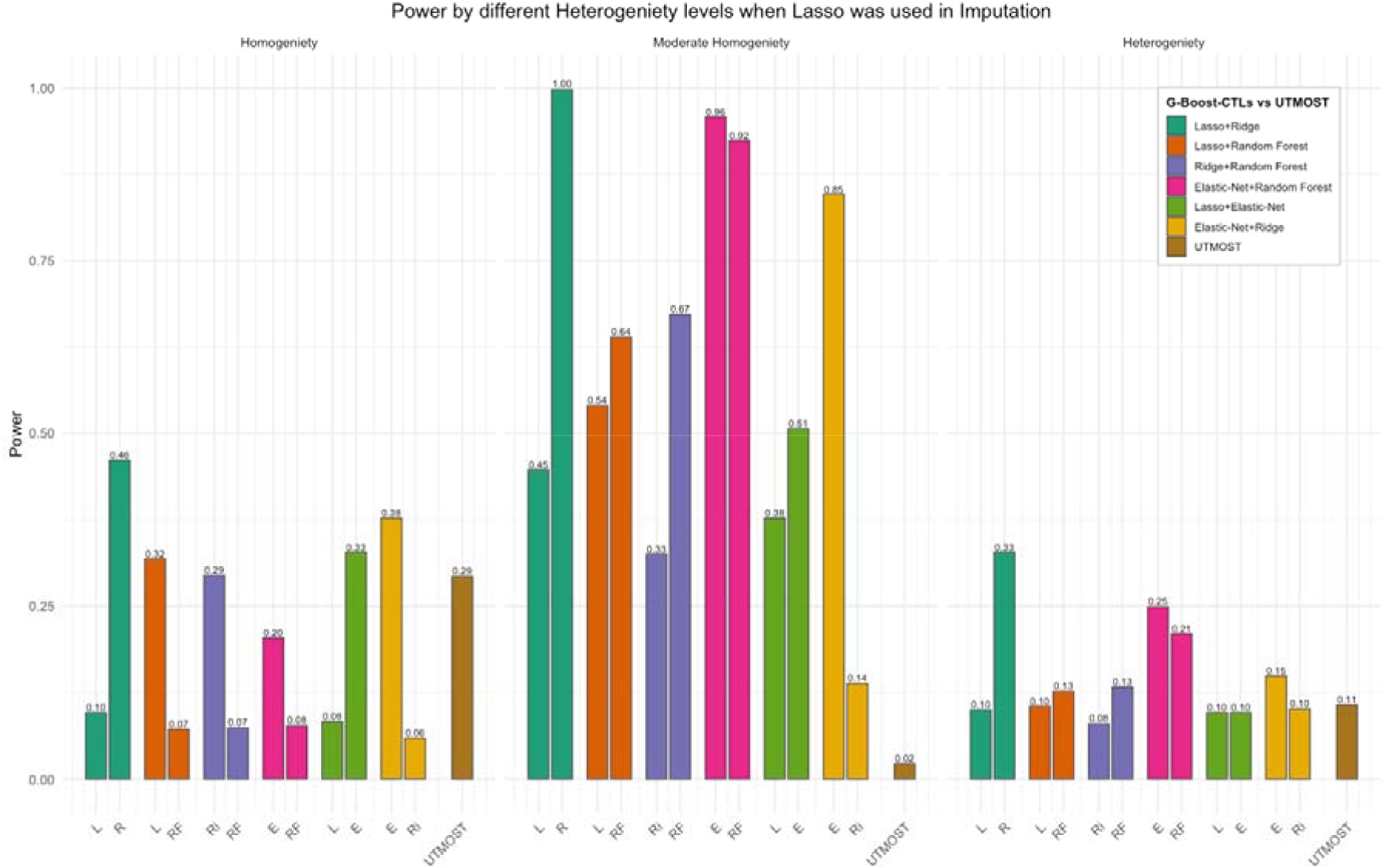
Statistical power comparison across varying levels of heterogeneity when Lasso is used for imputation in GBoost-CTL and UTMOST frameworks. Each panel corresponds to a different heterogeneity level (Homogeneity, Moderate Homogeneity, and Heterogeneity). Within each panel, the x-axis indicates which single-tissue learner (STL) was used to integrate GWAS information into the G-Boost-CTLs.

**Table 4.** Power Comparison of different GBoost-CTLs with UTMOST in different U settings (sample size 750, replication 10000)–when Lasso was used to impute 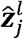. The highest power across different heterogeneity levels is bold.

| $U$ | L+Ri (L) | L+Ri (Ri) | L+RF (L) | L+RF (RF) | R+RF (Ri) | R+RF (RF) | E+RF (E) | E+RF (RF) | L+E (L) | L+E (E) | E+Ri (E) | E+Ri (Ri) | UT |
| --- | --- | --- | --- | --- | --- | --- | --- | --- | --- | --- | --- | --- | --- |
| **1** | 0.096 | **0.461** | 0.318 | 0.072 | 0.294 | 0.074 | 0.204 | 0.077 | 0.083 | 0.328 | 0.377 | 0.059 | 0.293 |
| **2** | 0.447 | **0.998** | 0.540 | 0.639 | 0.325 | 0.672 | 0.958 | 0.924 | 0.377 | 0.506 | 0.846 | 0.138 | 0.022 |
| **3** | 0.100 | **0.328** | 0.105 | 0.127 | 0.08 | 0.133 | 0.249 | 0.210 | 0.096 | 0.096 | 0.149 | 0.101 | 0.107 |

Table 5 and **Figure 6** illustrates the power comparison between various GBoost-CTL methods and UTMOST, based on gene expression imputation using UTMOST. In the homogeneous setting (U = 1), UTMOST exhibited the highest power (0.984), outperforming all G-Boost-CTLs, which remained below 0.19 - highlighting UTMOST’s strength in single-tissue dominant scenarios. For moderate heterogeneity (U = 2), while UTMOST achieved perfect power (1.000), Elastic Net–boosted CTLs like L + E (E) and E + RF (RF) performed competitively, reaching powers above 0.90. Under high heterogeneity (U = 3), UTMOST again dominated (0.998), but L + E (E) and E + RF (RF) remained the best-performing GBoost-CTLs (0.254 and 0.228). Across all U configurations, Elastic Net consistently emerged as the most effective boosting learner, outperforming Ridge- and Lasso-based variants. While UTMOST remains the overall top performer when used end-to-end, G-Boost-CTLs - especially those enhanced with Elastic Net-driven GWAS weights - provide competitive alternatives in heterogeneous environments, suggesting potential in hybrid frameworks that leverage UTMOST’s imputation with adaptive, interpretable weighting.

**Figure 6.**
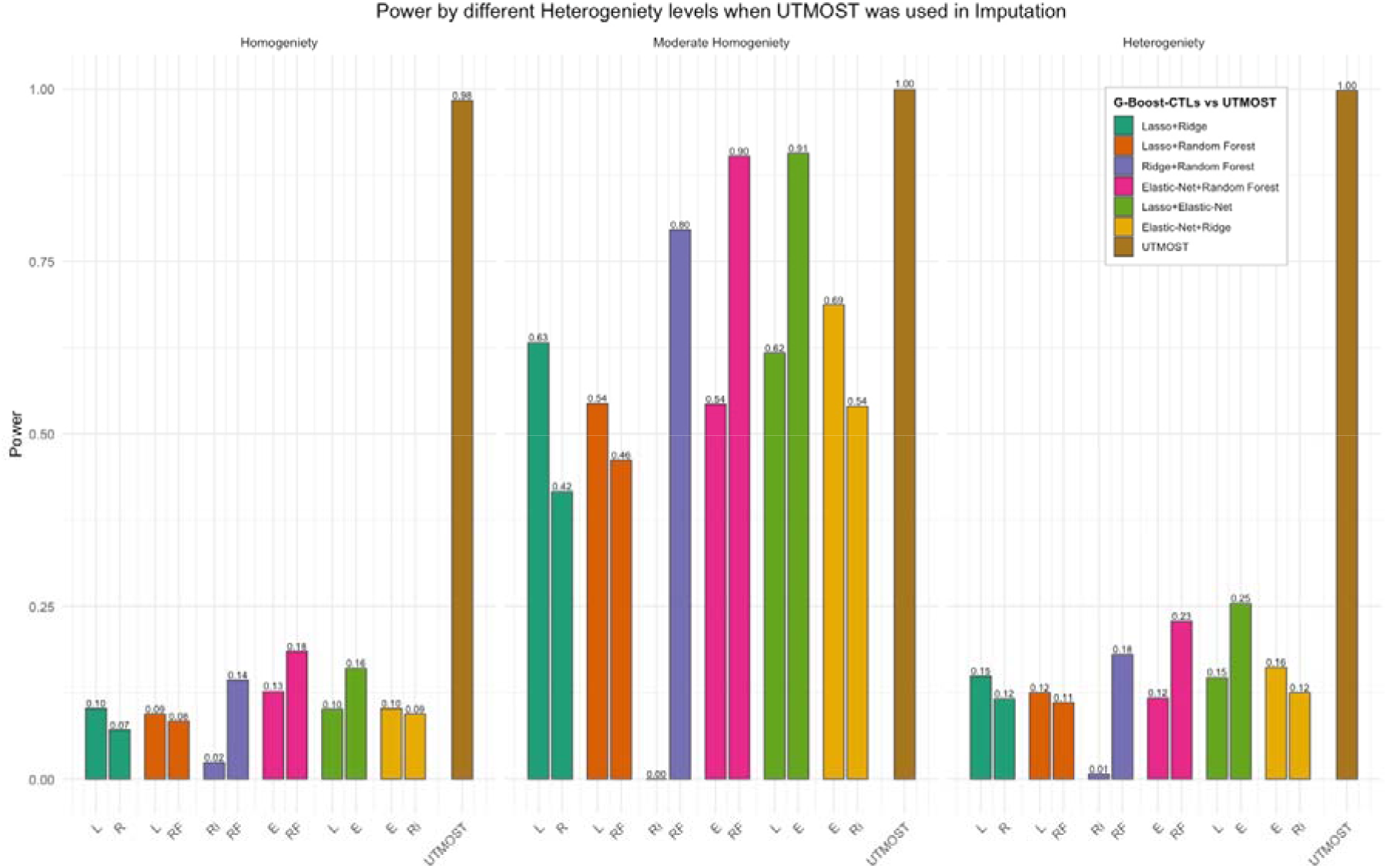
Statistical power comparison across varying levels of heterogeneity when UTMOST is used for imputation in GBoost-CTL and UTMOST frameworks. Each panel corresponds to a different heterogeneity level (Homogeneity, Moderate Homogeneity, and Heterogeneity). Within each panel, the x-axis indicates which single-tissue learner (STL) was used to integrate GWAS information into the G-Boost-CTLs.

**Table 5.** Power Comparison of different GBoost-CTLs with UTMOST in different U settings (sample size 750, replication 10000)–when UTMOST was used to impute 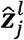. The highest power across different heterogeneity levels is bold.

| $U$ | L+Ri (L) | L+Ri (Ri) | L+RF (L) | L+RF (RF) | R+RF (Ri) | R+RF (RF) | E+RF (E) | E+RF (RF) | L+E (L) | L+E (E) | E+Ri (E) | E+Ri (Ri) | UT |
| --- | --- | --- | --- | --- | --- | --- | --- | --- | --- | --- | --- | --- | --- |
| 1 | 0.102 | 0.071 | 0.094 | 0.084 | 0.024 | 0.143 | 0.127 | 0.185 | 0.101 | 0.160 | 0.102 | 0.094 | **0.984** |
| 2 | 0.632 | 0.416 | 0.544 | 0.462 | 0.000 | 0.796 | 0.543 | 0.903 | 0.617 | 0.907 | 0.687 | 0.540 | **1.000** |
| 3 | 0.149 | 0.116 | 0.125 | 0.111 | 0.007 | 0.180 | 0.117 | 0.228 | 0.147 | 0.254 | 0.162 | 0.125 | **0.998** |

Finally, **Table 2.6** and **Figure 2.6** show the power comparison when the imputation was done with Random Forest. In this setting, where one tissue dominated the genetic influence on the phenotype, several GBoost-CTLs clearly outperformed UTMOST. The top-performing method was E+RF (E) with a power of 0.921, followed by L + Ri (R) at 0.786, and Ri + RF (Ri) at 0.762. UTMOST trailed at 0.106. These results reinforce the effectiveness of ensemble CTLs in prioritizing tissue-specific signals even when only one tissue contributed. Moderate homogeneity is the most favorable scenario for G-Boost-CTLs. Many methods, including Ri + RF (RF), E + RF (E), and E + RF (RF), achieved perfect or near-perfect power (1.000). Even combinations like L + RF (L) and L + RF (RF) scored 0.999, dramatically outperforming UTMOST (0.999) only marginally. However, UTMOST performed well here, reflecting that its design handles moderately heterogeneous tissue influences effectively. Still, G-Boost-CTLs delivered competitive or superior results by leveraging additional GWAS regulatory information. In heterogeneous situation, G-Boost-CTLs utilizing RF-based boosting, such as E + RF (RF) and Ri + RF (RF), maintained perfect power (1.000), while E + RF (E) achieved 0.998. UTMOST registered 0.579, a notable drop compared to the top-performing CTLs. These results indicate that G-Boost-CTLs are capable of maintaining high power even when the signal was equally distributed across diverse tissues—a setting typically challenging for traditional TWAS methods. A noteworthy observation is that RF as a GWAS weight learner (in combinations like L + RF (RF), Ri + RF (RF), and E + RF (RF)) generally yielded higher or comparable power compared to using Lasso, Ridge, or Elastic Net as the boosting learner. For instances, E + RF (RF) showed high power across all U settings: 0.439 (U=1), 1.000 (U=2), and 1.000 (U=3). Comparatively, L+RF (L) underperformed with the power of 0.612, 0.999, and 0.398 across the same settings. This suggests that Random Forest, being a non-linear and variance-aware model, is particularly effective in capturing the regulatory effects in GWAS-boosted frameworks.

**Table 6.** Power Comparison of different GBoost-CTLs with UTMOST in different U settings (sample size 750, replication 10000)–when Random Forest was used to impute 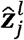. The highest power across different heterogeneity levels is bold.

| $U$ | L+Ri (L) | L+Ri (R) | L+RF (L) | L+RF (RF) | R+RF (Ri) | R+RF (RF) | E+RF (E) | E+RF (RF) | L+E (L) | L+E (E) | E+Ri (E) | E+Ri (Ri) | UT |
| --- | --- | --- | --- | --- | --- | --- | --- | --- | --- | --- | --- | --- | --- |
| **1** | 0.111 | 0.786 | 0.612 | 0.612 | 0.762 | 0.423 | **0.921** | 0.439 | 0.077 | 0.577 | 0.703 | 0.041 | 0.106 |
| **2** | 0.119 | 0.965 | 0.999 | 0.999 | **1.000** | **1.000** | **1.000** | **1.000** | 0.471 | 0.850 | 0.962 | 0.053 | 0.999 |
| **3** | 0.063 | 0.254 | 0.398 | 0.398 | 0.917 | **1.000** | 0.998 | **1.000** | 0.119 | 0.195 | 0.264 | 0.048 | 0.579 |

### Application to Real Data

To illustrate the effectiveness of our method, we applied it to the GOKIND dataset, which includes genotype and phenotype data relevant to type 1 diabetes. Following quality control, the dataset comprised 8,859 genes and 62,701 SNPs across 1,108 individuals. While the Bonferroni

correction yields a genome-wide threshold of approximately 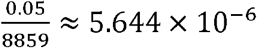, it is known to be overly conservative in multi-tissue TWAS due to its assumption of test independence. Therefore, following prior studies (Billah et al., 2025; Fischer et al., 2018), we adopted a more lenient significance level of 3 ×10^−4^ to strike a balance between discovery and false-positive control, particularly given the modest sample size of the GOKIND cohort relative to large-scale GWAS.

**Table 7** presents the gene-level findings from our G-Boost-CTL analysis. Trait annotations were drawn from the GWASL Catalog (https://www.ebi.ac.uk/gwas/home), and the “Description” field briefly summarizes previously reported associations, thereby placing each hit in a biological context. At the stringent Bonferroni threshold, GBoost-CTL pinpointed two genes: (i) FCHSD2 (11:72836745-73142318, p-value: 4.09 × 10^−7^), a locus linked to Systemic lupus erythematosus (Su et al., 2023), Crohn’s disease (Chranioti et al., 2022), Psoriasis (Abramczyk et al., 2020), HbA1c levels (Patiño-Fernández et al., 2009) - highlighting its association with type-1 diabetes (Billah et al., 2025); (ii) BFSP2 (3:133400056-133475222, p-value: 3.17 × 10^−6^), with its link with the traits like diffuse plaque measurement (Orchard et al., 2006), Serum Alanine Aminotransferase measurement (West et al., 2006), Platelet count (Schneider, 2009), Hemoglobin A1c (HbA1c) testing (Eyth & Naik, 2023), and Total Cholesterol measurement (Candler et al., 2017) - associated with type I diabetes (Billah et al., 2025).

**Table 7.**
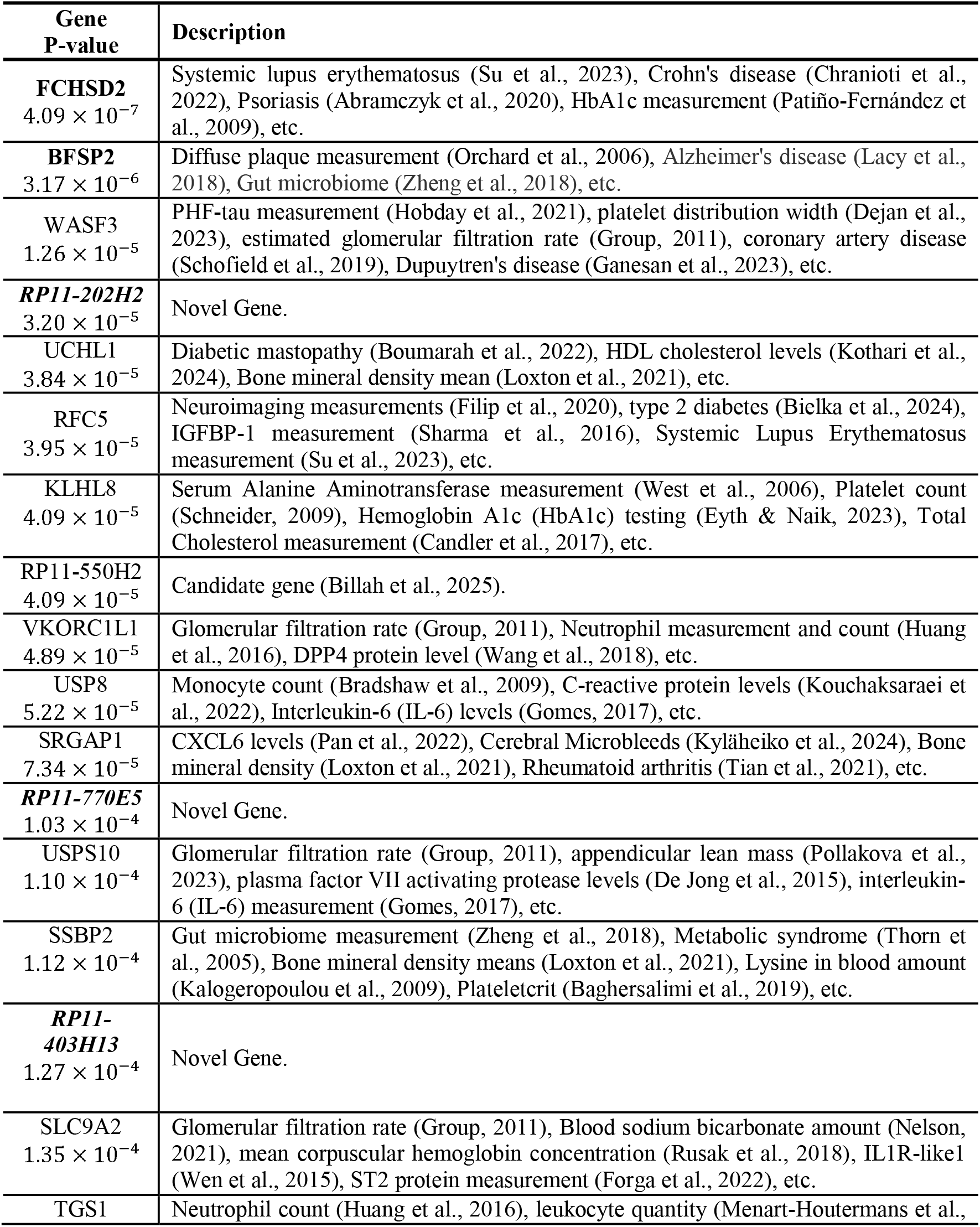
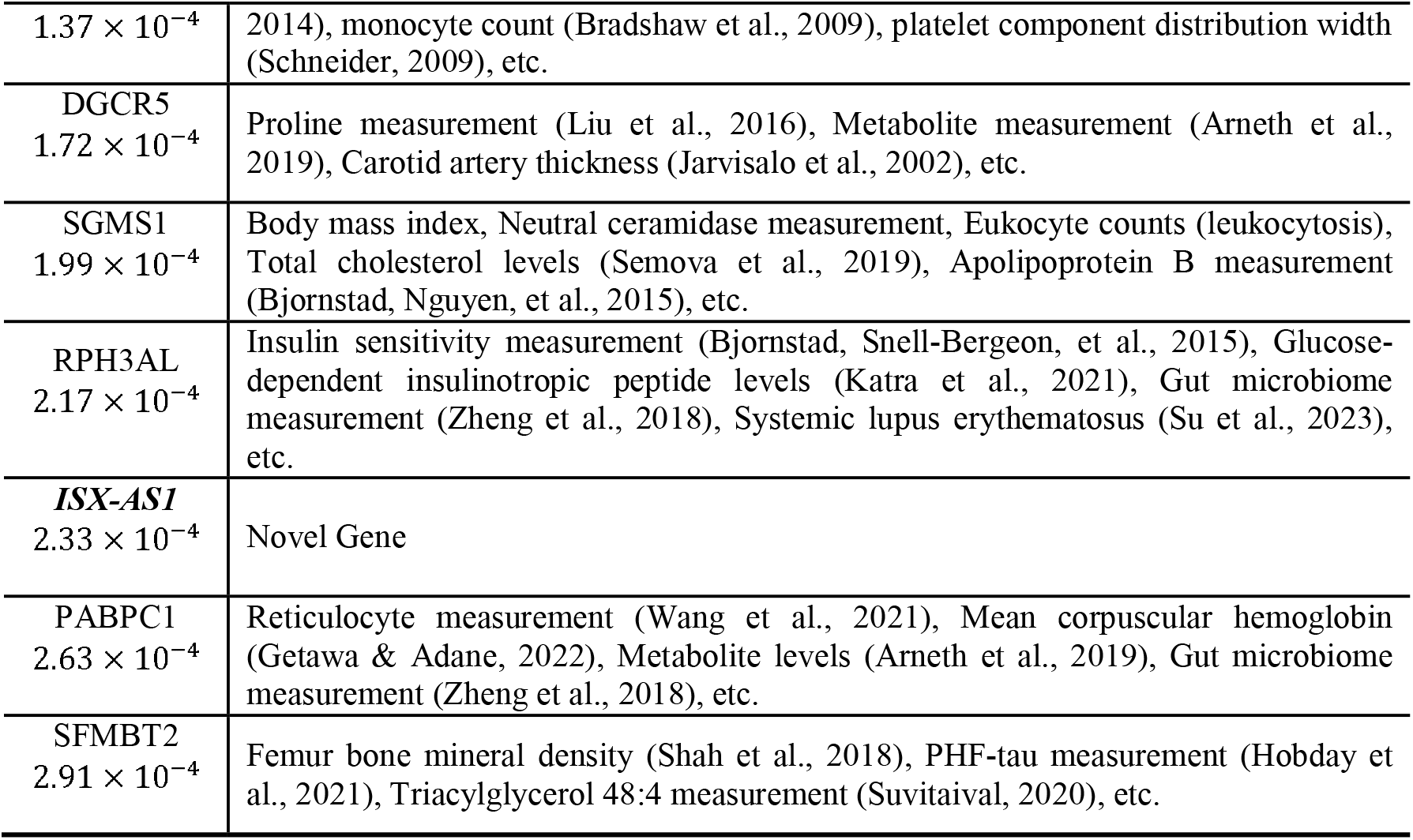
Genes significantly associated by GBoost-CTL at a p-value threshold of 3 × 10^−4^. Trait information in the “Description” column was sourced from the GWAS Catalog and reflects phenotypes previously linked to these genes. Citations indicate prior studies connecting the traits to type 1 diabetes. Genes with Bonferroni corrected threshold setting are bold and novel genes are bold and italic.

The p-values of these genes identified using GBoost-CTL are notably smaller than those reported under TWAS-CTL. For instance, the gene FCHSD2 exhibited a p-value of 2.81 × 10^−6^ under TWAS-CTL, whereas its significance under GBoost-CTL was even stronger. Similarly, BFSP2 had a p-value of 8.72 × 10^−6^ with TWAS-CTL, but was detected with greater confidence using G-Boost-CTL. Importantly, the previously highlighted novel gene - RP11-550H2, a long intergenic non-protein coding RNA (chr1:76,507,376-76,531,913) - was also identified by both methods. However, GBoost-CTL again yielded a smaller p-value of 4.09 × 10^−5^, compared to 2.90 × 10^−4^ under TWAS-CTL. This consistent improvement in statistical strength highlights the enhanced sensitivity of GBoost-CTL.

Beyond these replicated loci, the GBoost-CTL also identified four previously unreported Long Intergenic Non-Protein Coding RNAs to be associated with type I diabetes: (i) RP11-202H2 (chr12:86,993,355-87,232,775, p-value: 3.20 × 10^−5^), (ii) RP11-770E5 (chr8:48,658,981-48,696,887, p-value: 1.03 × 10^−4^), (iii) RP11-403H13 (chr9:6,902,670-6,978,859, p-value: 1.27 × 10^−4^), and (iv) ISX-AS1 (chr822:34,922,609-35,072,889, p-value: 2.33 × 10^−4^), and suggesting possible new regulatory elements worthy of functional follow-up and validation in independent cohorts.

Furthermore, in comparison to TWAS-CTL - which identified five genes at the *X*× 10^−5^ significance threshold - GBoost-CTL detected nine genes at the same level. At the study-wide suggested threshold of 3 × 10^−4^, G-Boost-CTL identified a total of 23 genes, outperforming TWAS-CTL, which discovered 15 genes; with an overlap of nine genes in between these methods (**Figure 8**). Notably, GBoost-CTL consistently uncovered a broader spectrum of associations even at stringent significance levels. In contrast, UTMOST identified only two genes - MAST2 (p-value: 2 × 10^−5^) and LINC00575 - while PrediXcan, combined with the GBJ test, identified one gene (KLHL8) with a p-value of 9.37 ×0 10^−5^. Interestingly, GBoost-CTL also identifies KLHL8, however, with a more stringent p-value of 4.09 × 10^−5^. When the p-value threshold was relaxed to 1 × 10^−3^, GBoost-CTL identifies 50 genes, whereas TWAS-CTL, UTMOST, and PrediXcan only detected 35, 4, and 8 genes, respectively. These comparisons underscore the enhanced sensitivity and broader discovery potential of GBoost-CTL, which is capable of identifying a larger set of biologically meaningful loci without compromising statistical rigor.

**Figure 7.**
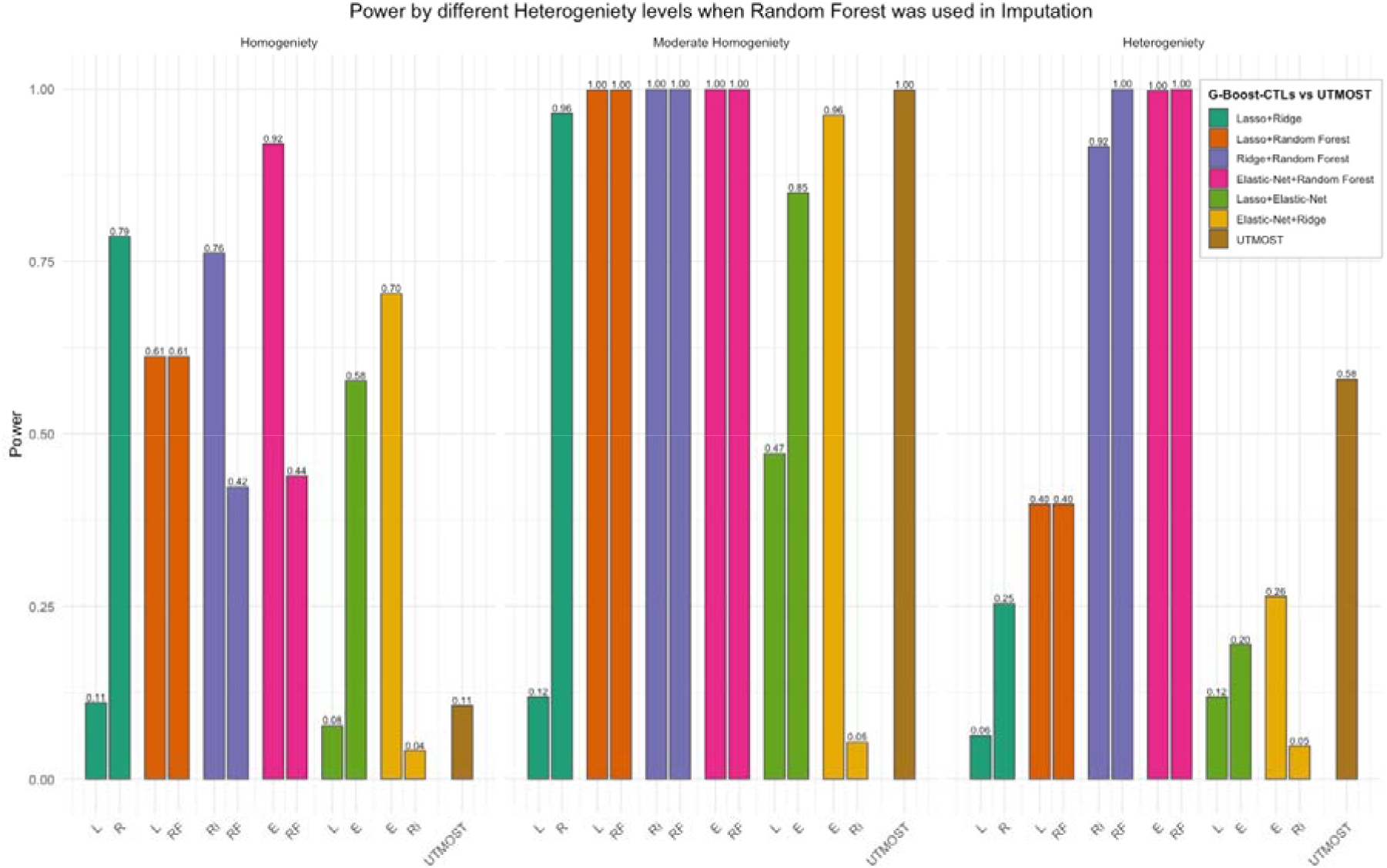
Statistical power comparison across varying levels of heterogeneity when Random Forest is used for imputation in GBoost-CTL and UTMOST frameworks. Each panel corresponds to a different heterogeneity level (Homogeneity, Moderate Homogeneity, and Heterogeneity). Within each panel, the x-axis indicates which single-tissue learner (STL) was used to integrate GWAS information into the G-Boost-CTLs.

**Figure 8.**
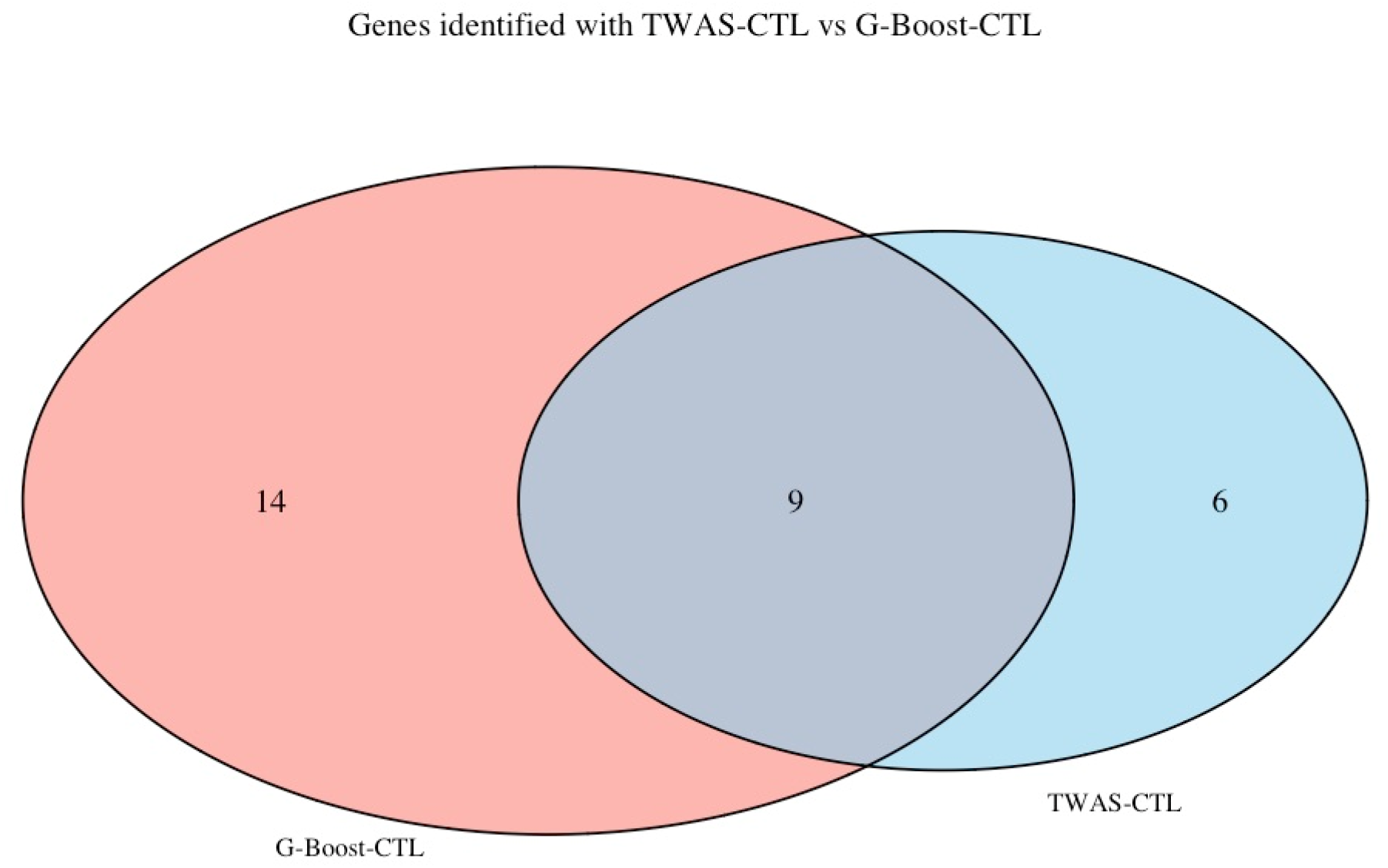
Comparison of TWAS-CTL and GBoost-CTL in identifying genes associated with TypeL1LDiabetes.

## DISCUSSION

This study introduces a novel mutli-tisue ensemble method in TWAS named G-Boost-CTL - which couples flexible single-tissue imputers with data-driven, GWAS-informed weighting - can outperform the prevailing linear and covariance-based frameworks used in transcriptome-wide association studies. Moreover, our framework can eclipse the performance of TWAS-CTL - the cross-tissue learner proposed by (Billah et al., 2025) - by replacing its “penalize-the-worst” utility weighting with a GWAS-informed scheme that elevates, rather than merely rescales, the most biologically informative tissues. Across an extensive grid of simulations and a real-data application to the GOKIND type 1 diabetes cohort, the superiority of GBoost-CTL is established.

GBoost-CTL maintained nominal type I error while delivering substantially higher power than UTMOST in almost every heterogeneity regime, when gene expression was imputed in statistical power simulation with any of the four learners we evaluated (Random Forest, PrediXcan, Ridge, or Lasso). This advantage was most striking under moderate tissue heterogeneity - the setting that best mirrors GTEx observations of shared-yet-unequal cis regulation - but it persisted, albeit at a smaller margin, when effects were either fully homogeneous or broadly dispersed. Two ingredients proved critical. First, the use of weighting function that prioritizes tissues with higher contribution can play a crucial role in identifying the intrinsic relationship between the eQTL effect on the genetic variants across the tissues–shared in withing and between human organs. Second, GWAS-derived information weights allowed GBoost-CTL to adapt to whichever tissue carried the informative signal in the target cohort, something UTMOST’s fixed covariance structure or TWAS-CTL’s weighting function cannot do. Notably, GBoost-CTLs boosted with Elastic Net or Random-Forest weights were the most stable across all scenarios, echoing theoretical work showing that mixed linear–non-linear ensembles can approximate a wide class of genetic architectures. The simulation patterns translated directly to the empirical setting.

Using a study-wide threshold of 3 × 10^−4^ - a compromise that balances discovery with control of false positives in moderately sized TWAS cohorts - GBoost-CTL identified 23 genes, versus 15 for TWAS-CTL, 2 for UTMOST, and a single locus for PrediXcan. Most of the loci recovered by the legacy methods were recaptured with more stringent p-values in most cases under G-Boost-CTL, and the framework also unearthed several long intergenic non-coding RNAs that have not yet appeared in the Type 1 diabetes literature.

Like all statistical frameworks, multi-tissue TWAS methodologies are not without their limitations. The proposed GBoost-CTL, while methodologically innovative, also introduces certain computational challenges - particularly in the context of training multiple learner combinations across 49 tissues. This level of complexity can be computationally demanding and may not scale efficiently without further optimization. Future methodological improvements could include tissue dimensionality reduction strategies or shared representation learning to compress information across similar tissues while retaining biological signal. In addition to this, in this current work we have used GWAS information directly from the raw genotype data, however, in most cases, only GWAS summary statistics data is available. In the future work, we plan to incorporate G-Boost-CTL in GWAS summary level data. Moreover, the promise of aggregation of linear and non-linear ensembles shows promising results and hence our future work will focus on exploring the application of advanced machine learning (e.g. gradient-boosted tree) and deep learning methods in the essence of multi-tissue TWAS framework. Furthermore, while the GOKIND dataset provides a valuable testbed for methodological development, its sample size is relatively small compared to modern biobank-scale resources. Moving forward, the application of GBoost-CTL to large-scale, ancestrally diverse cohorts such as the UK Biobank will be essential. These datasets offer the statistical power and heterogeneity necessary to validate novel associations and build high-resolution, tissue-specific regulatory maps, ultimately enhancing the biological interpretability and generalizability of multi-tissue TWAS findings.

Nonetheless, GBoost-CTL unites the predictive strength of modern machine-learning imputers with a principled, GWAS-informed weighting scheme, delivering a demonstrable uplift in gene-trait discovery over TWAS-CTL, UTMOST, and PrediXcan without sacrificing error control. Taken the extensive simulation and real-data analysis into consideration, GBoost-CTL reinforces that: (i) Adaptive weighting matters: Borrowing information from GWAS to modulate tissue weights complements the genetic-covariance strategy of UTMOST and reduces the risk of down-weighting the causal tissue when effects are unbalanced, (ii) Non-linearity is beneficial: Tree-boosted methods (e.g. Bootstrap aggregation like Random Forest) and Elastic-Net-weighted ensembles capture interaction and dominance effects that linear models systematically may underfit, and (iii) Statistical integrity is preserved: Despite the added flexibility, type I error remained at the nominal level across all sample sizes and replication depths, addressing long-standing concerns that complex learners might over-fit reference panels. As multi-tissue resources enlarge and computational barriers fall, such hybrid strategies like G-Boost-CTL are poised to become the new standard for elucidating the regulatory landscape of complex disease genetics.

